# Metabotropic group II glutamate receptors in the basolateral amygdala mediate cue-triggered increases in incentive motivation

**DOI:** 10.1101/2021.04.21.440686

**Authors:** Caroline Garceau, Anne-Noël Samaha, Thomas Cordahi, Alice Servonnet, Shaun Yon-Seng Khoo

## Abstract

**Rationale:** Reward-associated cues can trigger incentive motivation for reward and invigorate reward-seeking behaviour via Pavlovian-to-instrumental transfer (PIT). Glutamate signaling within the basolateral amygdala (BLA) modulates cue-triggered increases in incentive motivation. However, the role of BLA metabotropic group II glutamate (mGlu_2/3_) receptors is largely unknown.

**Objectives:** In Experiment 1, we characterized cue-triggered increases in incentive motivation for water reward using the PIT paradigm. In Experiment 2, we assessed the influence of intra-BLA microinjections of the mGlu_2/3_ receptor agonist LY379268 on this effect.

**Methods:** Water-restricted male Sprague-Dawley rats learned to press a lever for water. Separately, they learned to associate one of two auditory cues with free water. On test days, rats could lever press under extinction conditions (no water), with intermittent, non-contingent CS+ and CS- presentations. In Experiment 1, rats were tested under baseline conditions. In Experiment 2, rats received intra-BLA microinjections of LY379268 (0, 3 and 6 μg/hemisphere) before testing.

**Results:** Across experiments, CS+, but not CS- presentations increased water-associated lever pressing during testing, even though responding was reinforced neither by water nor the CS+. Intra-BLA LY379268 abolished both CS+ potentiated pressing on the water-associated lever and CS+ evoked conditioned approach to the site of water delivery. LY379268 did not influence locomotion or instrumental and Pavlovian response rates during intervals between CS presentations or during the CS-, indicating no motor effects.

**Conclusions:** mGlu_2/3_ receptor activity in the BLA mediates CS-triggered potentiation of incentive motivation for reward, suppressing both CS-induced increases in instrumental pursuit of the reward and anticipatory approach behaviour.

## Introduction

Environmental cues that predict rewards can acquire incentive motivational properties, enabling these cues to trigger and energize pursuit of the associated rewards (Bolles, 1972; Bindra, 1978). Cue-triggered increases in incentive motivation can be adaptive, guiding animals towards rewards needed for survival, such as food, water and safety. However, too much conditioned incentive motivation can contribute to excessive reward seeking as in addiction or overeating, and conversely too little of it can contribute to low levels of reward seeking, as in depression and anxiety (Olney et al., 2018; O’Brien et al., 1998; Ludwig et al., 1974; Everitt et al., 2001). It is therefore of interest to understand the neural mechanisms by which cues trigger incentive motivation to guide reward-seeking behaviour.

In both humans and laboratory animals, the amygdala and its basolateral complex (BLA) are involved in cue-triggered incentive motivation. For instance, activity in the amygdala increases during cue-evoked potentiation of reward seeking in humans (Talmi et al., 2008; Prevost et al., 2012). Furthermore, findings in laboratory animals show that neural activity in the BLA mediates the response to reward cues. First, neural activity in the BLA and its projections is necessary for the expression of cue-induced incentive motivation. Lesioning the BLA, inactivating CamKII-containing BLA neurons or inactivating BLA projections to the orbitofrontal cortex or the nucleus accumbens disrupt cue-triggered potentiation of reward seeking (Lichtenberg et al., 2017; Corbit & Balleine, 2005; Blundell et al., 2001; Derman et al., 2020; Shiflett & Balleine, 2010, see also Gabriele & See, 2010; Puaud et al., 2021). Similarly, pharmacological disconnection of the BLA and insular cortex disrupts CS-evoked conditioned approach behaviours (Nasser et al., 2018). Second, neural activity within the BLA is sufficient to potentiate the incentive motivational properties of reward cues. For instance, optogenetic stimulation of BLA neurons both potentiates cue-evoked expectation of the primary reward during Pavlovian conditioning and also invigorates instrumental responding for reward-associated cues (Servonnet et al., 2020). Within the BLA, glutamate-mediated transmission influences the behavioral response to reward-associated cues. Intra-BLA injections of agents that block AMPA (Malvaez et al., 2015), NMDA (Feltenstein & See, 2007) or metabotropic glutamate type 5 receptors (Khoo et al., 2019) disrupt cue-evoked appetitive responding.

In parallel, group II metabotropic glutamate receptors (mGlu_2/3_) are highly expressed in the BLA (Petralia et al., 1996; Ohishi et al., 1998; Gu et al., 2008), and they may also have a role in cue-triggered incentive motivation for reward. These receptors are localized predominantly extrasynaptically on presynaptic terminals, and their activation suppresses both synaptic glutamate release and excitability of projection neurons (Schoepp, 2001; Imre, 2007; Conn & Pin, 1997). Intra-BLA injections of an mGlu_2/3_ receptor agonist (LY379268) were found to have no effect on cue-induced reinstatement of extinguished cocaine-seeking behaviour (Lu et al., 2007), a response mediated at least in part by cue-triggered incentive motivation. This is in contrast to studies giving the agonist either systemically (Bossert et al., 2006; Baptista et al., 2004; Backstrom & Hyytia, 2005; Zhao et al., 2006) or into the central nucleus of the amygdala (Lu et al., 2007; Uejima et al., 2007), and showing reduced cue-induced reinstatement of extinguished reward-seeking behaviour. While these studies are informative, cue-induced reinstatement paradigms are not tests of pure cue-triggered incentive motivation, because instrumental responding at test is reinforced by cue presentation and thus also involves secondary reinforcement. As such, the contributions of BLA mGlu_2/3_ receptors to cue-triggered incentive motivation remain to be examined.

Here, we sought to determine the extent to which activation of BLA mGlu_2/3_ receptors modulates cue-triggered incentive motivation. To this end, we used Pavlovian-to-instrumental transfer (PIT). PIT is a test of pure cue-triggered incentive motivation, because it measures this process without the confounding influences of primary or secondary reinforcement (Cartoni et al., 2016; Walker, 1942; Rescorla & Solomon, 1967; Wyvell & Berridge, 2000). In PIT, subjects learn that a Pavlovian cue predicts a primary reward. Separately, they also learn to perform an instrumental response to obtain that reward. On test day, the subjects can perform the same instrumental response, but now under extinction conditions, where responding produces no reward and no cue. During testing, subjects receive free, intermittent presentations of the cue, and PIT is observed when cue presentation increases ongoing reward-seeking behaviour, indicating cue-triggered wanting of reward. Thus, in a first experiment, we characterized PIT under baseline conditions, because some studies in rats report that behavioural indices of PIT can be subtle (Wyvell & Berridge, 2000; Delamater & Holland, 2008). In a second experiment, we assessed the effects of intra-BLA microinjections of the mGlu_2/3_ agonist LY379268 immediately prior to PIT tests.

## Methods

### Animals

All procedures involving rats were *i*) approved by the animal ethics committee at the Université de Montréal, *ii*) followed ‘Principles of laboratory animal care’, and *iii*) adhered to Canadian Council on Animal Care guidelines. Adult male Sprague-Dawley rats (Charles River Laboratories, Montréal, Quebec, Canada) ordered at 200-225 g in Exp. 1 (N = 16) and 250-275 g in Exp. 2 (N = 16) on arrival, were housed 2 per cage in Exp. 1 or 1 per cage in Exp. 2, to avoid damage to intracerebral implants by conspecifics. The colony room was climate-controlled (22±1°C, 30±10% humidity) and maintained on a reverse 12-hour light/dark cycle (lights off at 8:30 a.m.). Experiments were conducted during the dark phase. Upon arrival, rats had free access to food (Rodent 5075, Charles River Laboratories) and water. After 72 hours of acclimation to the animal colony, rats were handled daily for at least 3 days. Beginning at 4 days after their arrival in Exp. 1 or 7 days after intracerebral surgery in Exp. 2, rats had restricted water access to facilitate acquisition of instrumental and Pavlovian conditioning, where water was the unconditioned stimulus (US). We used water as the US, because a water-paired cue has been shown to potentiate water-seeking behaviour in humans (i.e., a water-paired cue supports PIT) (De Tommaso et al., 2018). Rats first had 6 h/day of water access for 4 days, 4 h/day for the next 3 days, and then 2 h/day until the end of the experiments. Water was always given at least 1 hour after the end of testing, and at the same time each day.

### Behavioural Apparatus

Training and testing took place in standard operant chambers (31.8 x 25.4 x 26.7 cm; Med Associates, VT, USA) located in a testing room separate from the colony room. The operant chambers were placed in sound-attenuating boxes equipped with a fan that reduced external noise. Each chamber contained a tone generator located adjacent to the house light on the back wall, and a clicker located outside the chamber. On the opposite wall, there were two retractable levers (left active; right inactive) on either side of a water cup equipped with an infrared head entry detector. A liquid dispenser was calibrated to deliver 100 μL drops of water into the cup. Locomotor activity was measured using 4 infrared photobeams spaced evenly at floor level. Chambers were connected to a PC running Med-PC IV.

### Exp. 1: Characterizing Pavlovian-to-Instrumental Transfer

#### Instrumental conditioning

All training and testing procedures for general (non-selective) PIT were adapted from Derman & Ferrario (2018). Figure 1a illustrates the training and testing timeline. Rats (n = 16) first received instrumental conditioning where they learned to lever press for water. In the first session, water (100 μL) was available on a fixed-ratio 1 reinforcement schedule (FR1). A second inactive lever was present throughout, but had no programmed consequences. Sessions ended after rats earned 50 water deliveries or 40 min. Rats received FR1 sessions until they earned 50 water deliveries within a session, before transitioning to a variable interval (VI) reinforcement schedule. Using interval schedules of reinforcement during instrumental training subsequently promotes the expression of PIT (Lovibond, 1983). The VI schedule was increased across 8 sessions (40 min/session), in the following sequence: two VI10 sessions (range: 5-15 sec), two VI30 sessions (range: 15-45 sec) and four VI60 sessions (range: 30-90 sec). Lever pressing was recorded throughout each training session.

**Figure 1.**
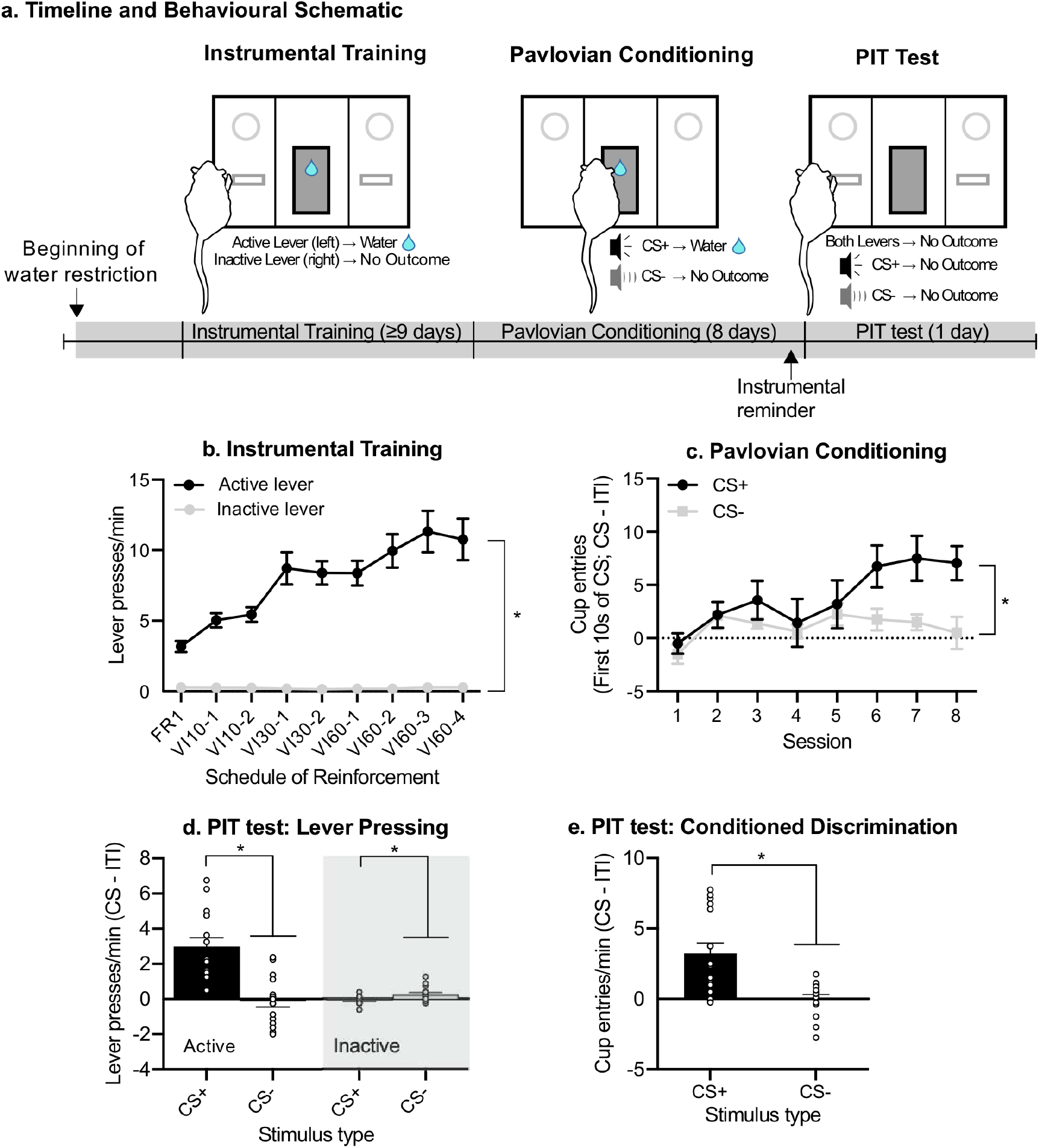
After instrumental and Pavlovian conditioning, rats show significant Pavlovian-to-Instrumental transfer. (a) Timeline and behavioural schematic for Experiment 1. (b) The rate of active lever presses increased over the daily 40-min instrumental training sessions. (c) The rate of water cup entries during the first 10 sec of each CS presentation was higher during CS+ relative to CS- presentations, and this difference increased over the daily 44-min Pavlovian conditioning sessions. (d) During a test for Pavlovian-to-instrumental transfer, CS+, but not CS- presentations invigorated lever pressing for water reward under extinction conditions. (e) During the test, rats also entered the water cup significantly more during CS+ compared to CS- presentations. In (c), water cup entries are shown as a difference score between responses during the first 10 sec of each CS presentation and during the last 10 sec of the inter-trial interval (ITI) immediately preceding each CS presentation. In (d-e), water cup entries/lever presses are shown as a difference score between responses during the 2-min CS and during the 2-min ITI immediately preceding each CS presentation. Data are presented as means ± SEM (N = 16). * *p* < 0.05. FR; fixed ratio. VI; variable interval. CS; conditioned stimulus.

#### Pavlovian conditioning

Next, rats received 8 Pavlovian conditioning sessions during which levers remained retracted, and rats now received presentations of two auditory stimuli, a 1800-Hz, 85-dB tone and a 10-Hz clicker (4 trials/CS). Each CS was presented for 2 min, because longer CSs (greater than 60 s) promote more robust PIT (Holmes et al., 2010). The mean inter-trial interval (ITI) was set at 180 s, ranging from 120-240 s, such that sessions were 44 min in total. One auditory stimulus was designated CS+ and concurrently paired with 4 US deliveries (100 μL water) on a VI30 schedule (range: 15-45 s; first water delivery ≥ 10 s from CS onset), while the other auditory stimulus (CS-) was also presented 4 times, but was not paired with water. CS+ and CS- were presented in alternation during each session. Rats were assigned to receive the tone-CS+ or clicker-CS+ such that mean active lever presses across the last four VI60 sessions and the number of FR1 sessions required to earn 50 water deliveries/session were similar across the groups. Water cup entries were recorded throughout each session to determine the extent to which rats learned the CS-US association.

#### Pavlovian-to-Instrumental transfer (PIT) testing

After Pavlovian conditioning, rats received an instrumental reminder session identical to VI60 training described above. PIT testing began on the next day. During each 42-min PIT session, both levers were available throughout. We measured lever-pressing behaviour under extinction conditions (no water was delivered) to assess cue-triggered incentive motivation. After 10 minutes, each CS (CS+ and CS-) was presented 4 times (2 min/presentation) in a counterbalanced order, with a fixed 2-min ITI. Lever responses, water cup entries and locomotion were recorded throughout the session.

### Exp. 2: Effects of activating mGlu_2/3_ receptors in the BLA on conditioned incentive motivation

#### Surgery

Figure 3a illustrates the experimental timeline. A separate cohort of rats (n = 16) weighing 325-375 g at the time of surgery was anesthetized with isoflurane (5% for induction, 2% for maintenance) and placed in a stereotaxic frame (Stoelting Co., IL, USA). Guide cannulae (26 GA, C315G, P1 Technologies, Roanoke, VA, USA) were implanted into the BLA at the following coordinates (in mm from bregma): AP −2.8, ML ± 5.0 and DV −6.2. Cannulae were occluded with dummies (C315CD) that were the same length as the cannulae. Injectors (33 GA, C315I, P1 Technologies) projected 2 mm beyond cannulae tips (final DV coordinate, −8.2 mm from Bregma). Four stainless steel screws were then anchored to the skull and cannulae were fixed with dental cement. Rats received preoperative carprofen (1.5 mg, s.c.) and penicillin (3000 IU, i.m.). Rats were given at least 7 days of recovery before beginning water restriction and behavioural training as described for Experiment 1.

#### Microinjection Procedures

After the instrumental reminder session, rats were habituated to the microinjection procedure with an intra-BLA microinjection of artificial cerebrospinal fluid (aCSF (1x); Cold Spring Harbor, 2011). Vehicle (0.5 μl aCSF) was infused over 1 min through injectors connected to 5-μL Hamilton syringes (7000 series, Hamilton Co., Reno, Nevada) controlled by a microsyringe pump (Harvard PhD 2000, Harvard Apparatus, Saint-Laurent, Canada). Injectors were left in place for 1 additional minute to allow diffusion. After this habituation microinjection, rats were returned to their home-cages.

Immediately prior to PIT tests, rats received bilateral infusions of aCSF, 3 or 6 μg/hemisphere of LY379268 (Cat No.: 2453, Batch No.: 9B/232416, CAS No.: 191471-52-0, Tocris Bioscience, Oakville, Ontario), in a counterbalanced, within-subjects design. Between PIT tests, rats received 2 instrumental sessions and 1 Pavlovian session.

### Histology

Following the last PIT test, rats were deeply anesthetised with urethane (1.2 g/kg, i.p.) and decapitated. Brains were dissected and sectioned into 40 μm-thick coronal slices at −20°C in a cryostat (Leica CM1850, Leica Biosystems, IL, USA). Coronal slices were then thaw-mounted on glass slides, stained with thionin and examined according to a rat brain atlas (Paxinos & Watson, 2007) by an experimenter blind to data. Six rats were excluded following histology, because at least one of the two cannulae (1/hemisphere) was not in the BLA. Two of these 6 rats had unilateral BLA placements, while the remainder had heterogenous placements dorsal, medial and/or lateral to the BLA. Behavioural data from these 6 rats are not shown due to heterogenous cannula placements and the lack of sufficient statistical power to demonstrate a significant PIT effect in this group.

### Statistical Analysis

Data were analysed using SPSS 26 (IBM, Armonk, NY, USA). In Exp. 1, two-way repeated measures (RM) ANOVA was used to analyse active vs. inactive lever-pressing across instrumental conditioning sessions (Lever × Session; both as within-subjects variables) and CS+ vs. CS- water cup entries across Pavlovian conditioning sessions (CS × Session; both as within-subjects variables). During PIT tests, number of lever presses during CS+ vs. CS- were analysed using a two-way RM ANOVA (Lever × CS; both as within-subjects variables). During PIT tests, number of water cup entries during CS+ vs. CS- were analysed using a two-tailed, paired t-test. We used Pearson’s correlation coefficient to analyse the relationship between the rats’ performance during Instrumental or Pavlovian conditioning and PIT testing and, because these analyses included CS+, CS- and ITI periods, unadjusted CS+, CS- and ITI response rates were used. In Exp. 2, two-way RM ANOVA was used to analyse Pavlovian (CS × Session; as within-subjects variables) and instrumental conditioning (Lever × Session; as within-subjects variables). Three-way RM ANOVA was also used to analyse LY379268 effects on lever pressing (LY379268 × CS × Lever; all as within-subjects variables). Two-way RM ANOVA was used to analyse LY379268 effects on water cup entries/locomotion during PIT tests (LY379268 × CS; both as within-subjects variables). When interaction and/or main effects were significant, effects were analysed further using Bonferroni-adjusted multiple post-hoc comparisons. Where Mauchly’s test of sphericity revealed a significant violation, the Greenhouse-Geisser correction was applied (for ε < 0.75). During PIT tests, lever-pressing rates during CS+ and CS- presentations, water cup entries and locomotion are presented as elevation scores over baseline responding, given by the 2-min ITI period immediately prior to each 2-min CS presentation (Corbit & Balleine, 2005; Shiflett & Balleine, 2010; Khoo et al., 2019; Derman & Ferrario, 2018). Data are expressed as mean ± SEM. The α level was set at < 0.05. Underlying data and Med-PC code will be available on Figshare (Garceau et al., 2021).

## Results

### Experiment 1. Characterization of Pavlovian-to-Instrumental Transfer

The rats increased their presses on the active lever across instrumental sessions (Fig. 1b; main effect of Session, F(2.16,32.34) = 15.29, *p* < 0.001, ε = 0.27). This is expected, as variable-interval schedules of reinforcement increase responding. In addition, across reinforcement schedules, rats pressed more on the active versus inactive lever (main effect of Lever, F(1,15) = 103.25, *p* < 0.001), and this difference increased across sessions (Lever × Session interaction, F(2.23,33.46) = 16.74, *p* < 0.001, ε = 0.28). Thus, the rats learned the action-outcome contingency.

Rats next received Pavlovian conditioning. Fig. 1c shows average rates of water cup entries during the first 10 sec of CS+ and CS- presentation (i.e., prior to US delivery). Rats visited the water cup progressively more across sessions (main effect of Session, F(7,105) = 4.57, *p* < 0.001), in particular during CS+ compared to CS- presentations (Fig. 1c; main effect of CS, F(1,15) = 6.93, *p* = 0.02; CS × Session interaction, F(7,105) = 2.81, *p* = 0.01). Thus, rats exhibited conditioned discrimination, indicating learning of the Pavlovian CS+/water association.

#### Cue-induced incentive motivation for water reward

During PIT tests, rats could lever press, but this was not reinforced by water or the CS+ water cue. First, across the PIT test session, rats pressed more on the active than on the inactive lever (Fig. 1d; main effect of Lever, F(1, 5) = 24.96, *p* < 0.001), indicating significant water-seeking behaviour under extinction conditions, when no water was delivered. Second, the rats pressed more on the active lever during CS+ versus CS- presentations (main effect of CS F(1,15) = 17.51; CS × Lever interaction, F(1,15) = 26.33; active lever pressing during CS+ > CS-; All *p*’s < 0.001). The rats also pressed less on the inactive lever during CS+ presentations than they did during CS- presentations (*p* = 0.03). Thus, the CS+ both increased pressing on the water-associated lever and narrowed behavioural responding by decreasing pressing on the lever not associated with water. In contrast, a control cue (CS-) had no effect on lever-pressing behaviour.

Finally, rats exhibited cue-triggered discrimination of conditioned approach, preferentially visiting the water cup during CS+ versus CS- presentations (Fig. 1e; Paired t-test; t(15) = 4.61, *p* < 0.001). Together, these results indicate significant CS-triggered incentive motivation for water reward, as shown by PIT.

#### Behavioural performance during instrumental and Pavlovian conditioning predicts later cue-induced incentive motivation

We examined whether responding during previous instrumental and Pavlovian conditioning predicted behaviour during tests for CS+ triggered incentive motivation. More active lever pressing during the final instrumental conditioning session (VI60-4 in Fig. 1b) predicted more active lever pressing at test, specifically during CS+ presentations (Fig. 2a; r(14) = 0.71, r^2^ = 0.50, *p* = 0.002). The level of active lever pressing during the final instrumental conditioning session did not significantly predict active lever pressing during either CS- presentations (Supplementary Fig. 1a; r(14) = 0.34, r^2^ = 0.12, *p* = 0.20) or during the ITI at test (Supplementary Fig. 1b; r(14) = 0.21, r^2^ = 0.05, *p* = 0.43). The level of active lever pressing during the final instrumental conditioning session also did not significantly predict water cup entries at test (Supplementary Fig. 1c; r(14) = 0.50, r^2^ = 0.25, *p* = 0.051; (Supplementary Fig. 1d; r(14) = 0.092, r^2^ = 0.01, *p* = 0.73; Supplementary Fig. 1e; r(14) = −0.23, r^2^ = 0.05, *p* = 0.40). Thus, rats more sensitive to water reinforcement during prior instrumental conditioning later showed greater cue-induced incentive motivation for water reward, as shown specifically by more CS+ triggered presses on the previously water-associated lever.

**Figure 2.**
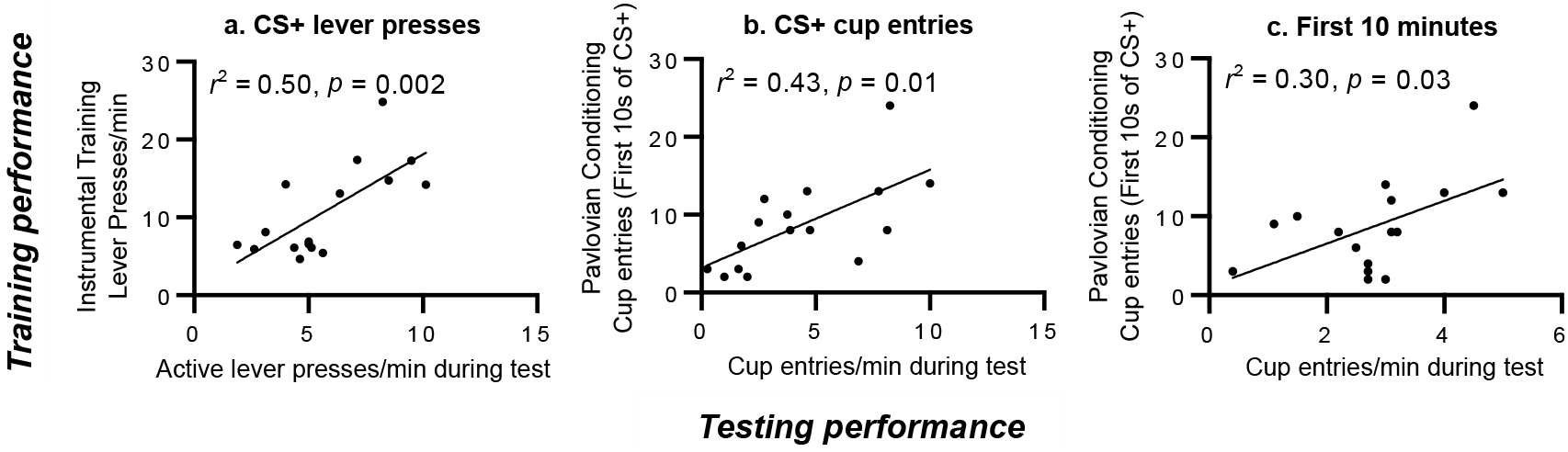
Responding during both instrumental and Pavlovian conditioning predicted later performance during a test for Pavlovian-to-instrumental Transfer. (a) More active lever pressing during the final instrumental training session predicted more active lever pressing at test, specifically during presentations of the water-paired conditioned stimulus (CS+). (b) More water cup entries during CS+ presentation on the last Pavlovian session (session 8) predicted more water cup entries at test during the CS+. (c) More water cup entries during CS+ presentation on the last Pavlovian session also predicted more water cup entries during the first 10 min of the PIT test session, when neither the CS+ nor water were presented (i.e., under extinction conditions). N = 16. Response rates during the CS are unadjusted for baseline (i.e., during the inter-trial interval, or ITI). CS; conditioned stimulus.

Performance during Pavlovian sessions also predicted responding at test. More CS+ elicited water cup entries during the final Pavlovian conditioning session (session 8 in Fig. 1c) significantly predicted more water cup entries at test, specifically during CS+ presentations (Fig. 2b; r(14) = 0.65, r^2^ = 0.43, *p* = 0.01). There were no significant correlations during CS- presentations (Supplementary Fig. 1f; r(14) = 0.30, r^2^ = 0.09, *p* = 0.27) or during the ITI at test (Supplementary Fig. 1g; r(14) = 0.06, r^2^ = 0.003, *p* = 0.83). Thus, more robust CS+ triggered conditioned approach behaviour during Pavlovian conditioning predicted a greater CS+ triggered conditioned approach response at test. More CS+ elicited water cup entries during the final Pavlovian conditioning session also significantly predicted more water cup entries during the first 10 min of the PIT test session, when neither the CS+ nor water were presented (Fig. 2c; r(14) = 0.55, r^2^ = 0.30, *p* = 0.03). Thus, rats that showed more CS+ triggered conditioned approach behaviour during Pavlovian conditioning also later visited the water cup more frequently under extinction conditions. Finally, CS+ elicited water cup entries during the final Pavlovian session did not significantly predict active lever pressing behaviour at test (Supplementary Fig. 1h, during CS+ presentations; r(14) = 0.18, r^2^ = 0.03; Supplementary Fig. 1i, during CS- presentations; r(14) = −0.40, r^2^ = 0.15; Supplementary Fig. 1j, during the ITI; r(14) = −0.41, r^2^ = 0.17; All *p*’s > 0.05). Thus, CS+ triggered approach behaviour during Pavlovian conditioning is dissociable from later instrumental pursuit of the primary reward.

### Experiment 2. Effects of activating mGlu_2/3_ receptors in the BLA on conditioned incentive motivation

Six rats did not have bilateral cannulae in the BLA, and they were excluded from data analysis. Over the course of instrumental conditioning, rats increased active lever pressing (Fig. 3b; main effect of Session, F(2.189,19.705) = 6.58, *p* = 0.005, ε = 0.27). In addition, rats pressed more on the active than on the inactive lever (main effect of Lever, F(1,9) = 54.71, *p* < 0.001) and this difference increased across sessions (Lever × Session interaction, F(2.128,19.153) = 6.94, *p* = 0.005, ε = 0.27). Thus, rats learned the action-outcome contingency, and reliably lever pressed for water.

**Figure 3.**
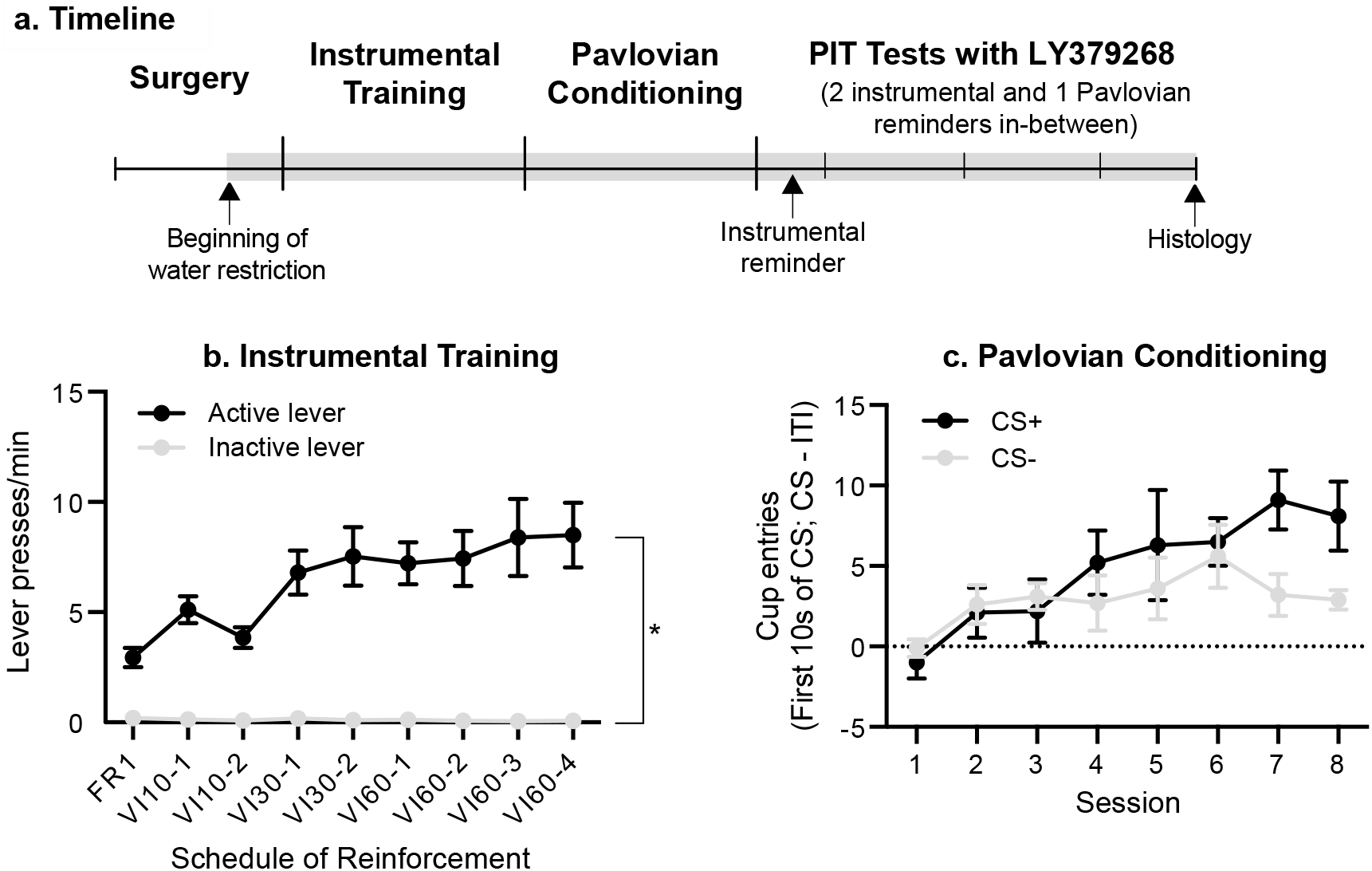
Timeline and acquisition of instrumental and Pavlovian conditioning in Experiment 2. (a) After implantation of bilateral cannulae targeting the basolateral amygdala (BLA), rats (N = 10) received instrumental and Pavlovian conditioning sessions. We then assessed the effects of LY379268 on Pavlovian-to-instrumental transfer. (b) The rate of active-lever presses increased over the daily 40-min instrumental training sessions. (c) The rate of water cup entries during the first 10 sec of each CS presentation increased over the daily 44-min Pavlovian conditioning sessions. Data are presented as means ± SEM. \**p* < 0.05. FR; fixed ratio. VI; variable interval. CS; conditioned stimulus. ITI; inter-trial interval.

Across Pavlovian conditioning sessions, rats increased their visits to the water cup during the first 10 sec of CS presentation (Fig. 3c; main effect of Session F(2.548,22.93) = 3.27, *p* = 0.046, ε = 0.36). CS+ water cup entries appeared higher than CS- water cup entries, though this was not statistically significant (main effect of CS, F(1,9) = 3.76, *p* = 0.08). There was also no significant change in conditioned discrimination across sessions (CS × Session interaction, F(7,63) = 1.77, *p* = 0.109).

#### Effects of intra-BLA LY379268 on lever pressing

Fig. 4a shows the placements of microinjector tips in the BLA (n = 10). Presentation of the CS+, but not the CS-, increased active lever pressing (Figs. 4b-c; main effect of CS, F(1, 9) = 9.4, *p* = 0.013; CS × Lever interaction (F(1, 9) = 8.61, *p* = 0.02; CS+ > CS-, *p* = 0.02), without affecting inactive lever pressing (*p* = 1). Thus, the CS+ selectively increased pressing on the water-associated lever—indicating PIT—without increasing general lever-pressing behaviour indiscriminately. LY379268 microinjection influenced lever-pressing behaviour, preferentially reducing active-versus inactive lever pressing (Figs. 4b-c; main effect of LY379268, F(2, 18) = 11.73, *p* < 0.001; LY379268 x Lever interaction, F(1.28, 11.48) = 12.73, *p* < 0.003, ε = 0.64). Compared to vehicle, 3 and 6 μg LY379268 reduced total active lever presses (i.e., active lever presses during both CS+ and CS- presentations. Fig. 4b; *p* = 0.003 and 0.042 respectively), with no significant effects on total inactive lever pressing (Fig. 4c; All *p*’s = 1). As Fig. 4c shows, pressing on the inactive lever was already low under baseline (intra-BLA vehicle) conditions, indicating a potential floor effect. Importantly, the effects of LY379268 on active lever pressing depended on CS type. LY379268 reduced active lever pressing during CS+, but not CS- presentations (Fig. 4b; CS × LY379268 interaction, F(2, 18) = 4.20, *p* = 0.03. Note that active lever pressing during CS- presentations was also already low at baseline). Compared to vehicle, 3 and 6 μg LY379268 reduced active lever presses during CS+ presentations (*p* = 0.002 and 0.03 respectively). Thus, LY379268 specifically attenuated CS+ triggered potentiation of water-seeking behaviour. This effect is further highlighted by analysing the influence of LY379268 on average PIT magnitude, indicated by the difference in active lever pressing during CS+ vs. CS- presentations. LY379268 reduced PIT magnitude (Fig. 4d; main effect of LY379268, F(2, 18) = 4.41, *p* = 0.03; 3 μg vs. vehicle, *p* = 0.03; 6 μg vs. vehicle, *p* = 0.07; 6 μg vs. 3 μg, *p* = 1). Thus, the CS+ enhanced responding on the water-associated lever, indicating cue-triggered potentiation of incentive motivation for water, and activating BLA mGlu_2/3_ receptors with LY379268 suppressed this effect.

**Figure 4.**
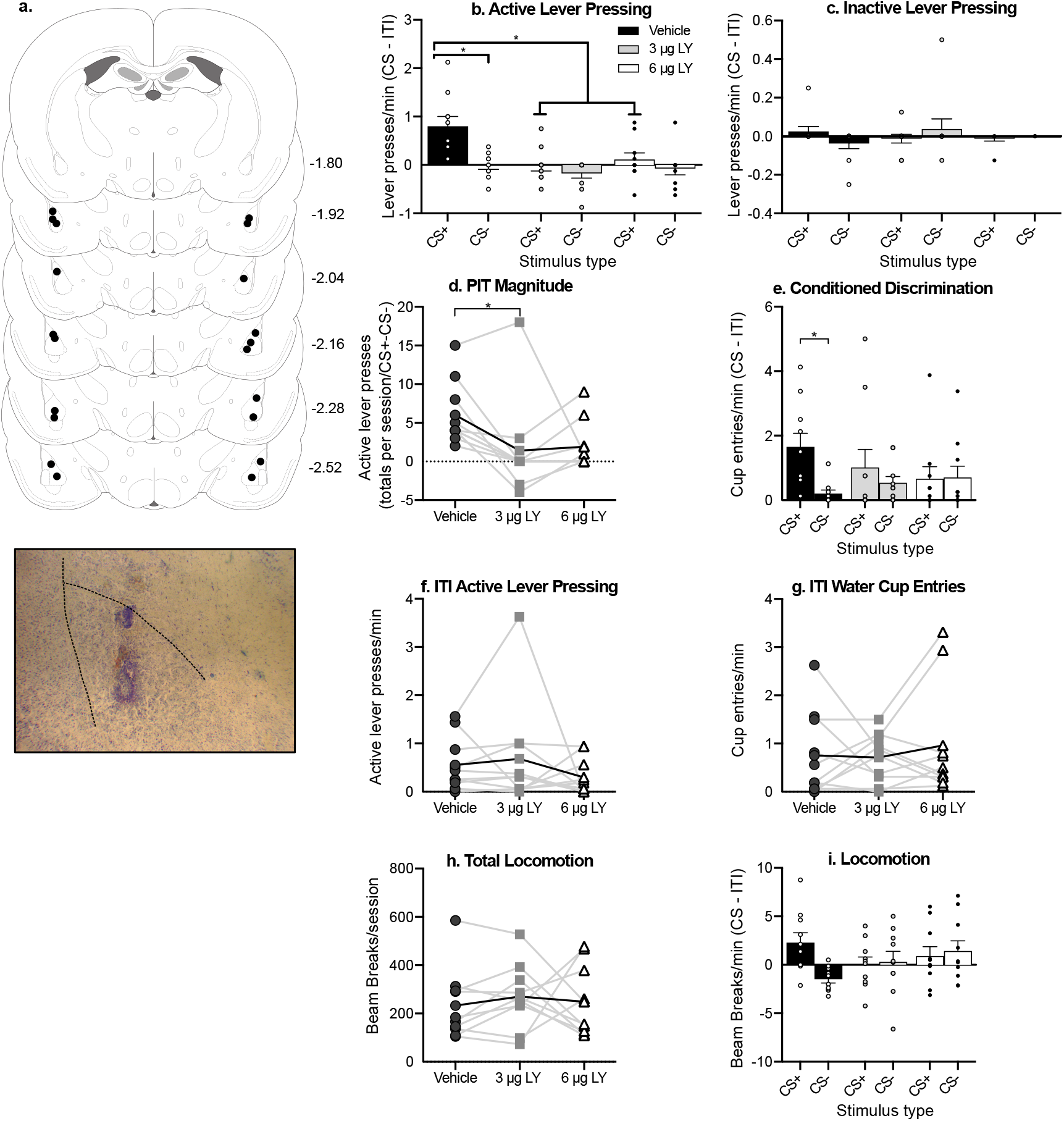
Activation of basolateral amygdala (BLA) mGlu2/3 receptors with LY379268 abolished CS+ triggered increases in both instrumental reward-seeking actions and conditioned approach behaviours. (a) Estimated placements of microinjector tips mapped to Rat brain atlas coordinates (Paxinos & Watson, 2007). An example photomicrograph is shown below. (b) At baseline (‘Vehicle’), presentation of the Pavlovian water cue (CS+) triggered increased responding on the water-associated lever compared to CS- presentation, and intra-BLA LY379268 (3 or 6 μg/hemisphere) abolished this effect. (c) LY379268 had no effect on inactive lever presses. (d) At 3 μg/hemisphere, LY379268 significantly reduced the magnitude of Pavlovian-to-instrumental transfer. (e) At baseline (‘Vehicle’), rats entered the water cup significantly more often during CS+ versus CS- presentations, and intra-BLA LY379268 (3 or 6 μg/hemisphere) abolished this Pavlovian conditioned approach behaviour. (f-g) LY379268 had no effect on active lever presses or water cup entries during inter-trial intervals (ITI). (h) LY379268 did not influence total locomotor activity, or (i) locomotor activity during CS presentations at test. In (d), and (f-h), thicker curve in each panel represents group means. Bar graphs present data as means ± SEM (n= 10). * *p* < 0.05. CS; conditioned stimulus.

Fig. 4e shows that under baseline conditions (‘vehicle’), the rats visited the water cup more frequently during CS+ versus CS- presentations, indicating significant conditioned discrimination (LY379268 × CS interaction, (F(2, 18) = 5.57, *p* = 0.01; under Vehicle, CS+ > CS-, *p* = 0.004). Intra-BLA LY379268 injections abolished this conditioned discrimination (All *p*’s > 0.05). Thus, the CS+ water cue triggered more visits to the water cup than the control cue did, and LY379268 prevented this conditioned discrimination effect.

Intra-BLA LY379268 did not produce significant motor suppressive effects. Compared to vehicle microinjections, LY379268 microinjections did not influence active lever pressing (Fig. 4f; F(1.26, 11.37) = 0.84, *p* = 0.41, ε = 0.63) or water cup entries during the ITI (Fig. 4g; F(2, 18) = 0.30, *p* = 0.75). LY379268 microinjections also had no influence on either total locomotor activity during testing (Fig. 4h; F(1.22,10.99) = 0.195, *p* = 0.72, ε = 0.61), or on locomotor activity during CS presentations (Fig. 4i; Main effect of Dose, F(2, 18) = 0.59, *p* = 0.56; Dose × CS interaction F(2, 18) = 3.27, *p* = 0.06). As such, the observed effects of intra-BLA LY379268 on cue-triggered potentiation of water-seeking behaviour cannot be attributed to motor suppressive effects.

## Discussion

Here we characterized the ability of a Pavlovian water cue to trigger increases in instrumental pursuit of water reward, when responding involved neither primary reinforcement by water nor conditioned reinforcement by the water cue. We then assessed the effects of intra-BLA microinjections of the mGlu_2/3_ receptor agonist LY379268 on this effect. Rats initially received instrumental conditioning sessions, where they learned to press a lever for water reward. The rats also received separate Pavlovian conditioning sessions, where they learned that a cue (CS+) predicts free water deliveries, and a second cue does not (CS-). We then used PIT procedures to assess cue-triggered increases in water-seeking behaviour where responding was reinforced neither by the cue nor by water reward. We report three key findings. First, during PIT testing, rats responded more for water during CS+ versus CS- presentations, indicating cue-triggered potentiation of incentive motivation for water reward. Second, responding during both the previous instrumental and Pavlovian conditioning phases significantly predicted later performance in the PIT test. Third, activation of mGlu_2/3_ receptors in the BLA impaired cue-triggered increases in reward seeking, without influencing general motor behaviour. Thus, increased mGlu_2/3_ receptor activity in the BLA attenuates the ability of reward-predictive cues to invigorate reward-seeking actions.

### Performance during previous instrumental and Pavlovian conditioning influences later cue-triggered responses

Performance during training selectively predicted behaviours triggered by the water-associated cue at test, demonstrating how acquired instrumental and Pavlovian associations shape later behaviour. First, responding during initial instrumental conditioning predicted behaviour at test. Higher rates of lever-pressing for water during instrumental conditioning predicted more robust instrumental responding for water reward at test, but only during presentations of the water-associated CS+. Performance during previous instrumental conditioning did not predict water-seeking behaviour during CS- presentations or during the time intervals in between CS presentations (ITI) at test. One possibility is that when rats attribute more reinforcing properties to a primary reward such as water, they will later show more vigorous cue-induced incentive motivation for that reward. In parallel, such rats might also be more susceptible to the response-*suppressing* effects of stimuli that do not predict the pursued reward (the CS- here). Indeed, CS+ triggered increases in reward-seeking actions may reflect both a facilitation of responding by the CS+ and a suppression of responding by the CS- (Holmes et al., 2010). Whatever the underlying mechanisms, our findings extend previous work showing that when instrumental training precedes Pavlovian conditioning—as used here—, more instrumental training predicts more cue-triggered reward pursuit later on (Holmes et al., 2010).

Performance during initial Pavlovian conditioning also predicted behaviour at test. More CS+ triggered water cup entries during Pavlovian conditioning predicted more CS+ triggered water cup visits at test. This effect was selective, as there were no such predictive relationships for water cup entries during CS- presentations or during the ITI at test. Because no water is delivered at test, our findings therefore suggest that rats in which a reward cue becomes a more important elicitor of approach responses could also be more resistant to extinction when the cue no longer predicts water. This provides a testable hypothesis to explore individual differences in conditioned appetitive responding.

### Activation of mGlu_2/3_ receptors in the BLA suppresses cue-triggered increases in incentive motivation: Behavioural specificity

Our findings support the idea that activation of BLA mGlu_2/3_ receptors reduces the expression of cue-triggered “wanting” for water, presumably by suppressing the incentive salience of the cue-evoked neural representation of water reward (see (Wyvell & Berridge, 2000). The mGlu_2/3_ receptor agonist, LY379268 abolished cue-triggered potentiation of incentive motivation for reward, as measured by *i*) the rate of pressing on the water-associated lever when the CS+ cue is presented, *ii*) PIT magnitude (the difference between active lever pressing during CS+ vs CS- presentation), and *iii*) CS+ triggered conditioned approach behaviour to the site of water delivery.

The suppressive effects of intra-BLA LY379268 injections on cue-induced increases in water cup entries could also reflect attenuation of the cue’s predictive value. Indeed, cue-elicited water cup entries are goal-tracking responses, and such responses are thought to preferentially reflect a cue’s predictive value (Flagel et al., 2007; Robinson & Flagel, 2009). In parallel, sign-tracking behaviours (i.e., where rats approach and interact with reward-associated cues) are thought to preferentially reflect a cue’s incentive motivational value Berridge & Robinson (2003); Flagel et al. (2011). However, the present study was not designed to distinguish goal-versus sign-tracking responses. Indeed, we used an auditory conditioned cue. Such cues are non-localizable, and are therefore less amenable to approach-like, sign-tracking behaviours. In comparison, studies designed to detect goal-versus sign-tracking phenotypes generally use tactile, localizable cues. This being said, the Pavlovian-to-Instrumental transfer paradigm we used permits simultaneous measurement of cue-induced Pavlovian and instrumental responses, thus allowing us to study the effects of intra-BLA LY379268 injections on both types of conditioned behaviours.

Intra-BLA injections of LY379268 influenced the behavioural response to the CS+ without producing significant motor-suppressing effects. Indeed, LY379268 had no effect on lever pressing behaviour or water cup entries during inter-trial intervals (i.e., in the absence of the CS+), or on locomotor activity. This is consistent with previous findings showing that at doses similar to those used here, intra-BLA LY379268 injections do not impair motor behaviour (Cannady et al., 2011). Thus, together our findings suggest that activating BLA mGlu_2/3_ receptors, which can suppress both local synaptic glutamate release and excitability of BLA projection neurons (Schoepp, 2001; Imre, 2007; Conn & Pin, 1997), decreases the incentive salience of reward-associated stimuli, causing animals to reduce their instrumental pursuit of the reward in a pure conditioned incentive paradigm.

The question now is how activation of BLA mGlu_2/3_ receptors suppresses cue-triggered potentiation of reward seeking actions. When activated, mGlu_2/3_ receptors provide negative feedback to decrease evoked glutamate release (Schoepp, 2001; Imre, 2007; Conn & Pin, 1997). We did not measure extracellular glutamate here. However, we hypothesize that synaptic glutamate is necessary for cue-triggered increases in incentive motivation, and that mGlu_2/3_ activation suppresses this conditioned effect by reducing synaptic glutamate release. This interpretation would be in agreement with Malvaez et al. (2015), who found that BLA glutamate release frequency correlates with lever-pressing actions that are selectively invigorated by a reward-associated CS, and that blockade of BLA AMPA glutamate receptors disrupts the conditioned incentive impact of that CS.

Malvaez et al. (2015) found that intra-BLA AMPA receptor blockade suppressed CS-triggered increases in the instrumental pursuit of reward, but that neither intra-BLA AMPA nor NMDA receptor blockade changed CS-triggered Pavlovian conditioned approach behaviour. We further extend these findings by showing that activation of intra-BLA mGlu_2/3_ receptors attenuates CS-triggered Pavlovian conditioned approach behaviour. In contrast to AMPA and NMDA receptor blockade, activating mGlu_2/3_ receptors suppresses synaptic glutamate release, thus presumably decreasing signaling at all local glutamate receptors, both metabotropic and ionotropic. Thus, our findings combined with those of Malvaez et al. (2015) suggest that activity at non-ionotropic glutamate receptors in the BLA mediate conditioned approach to reward-associated cues. This could involve metabotropic glutamate type 5 receptors (Khoo et al., 2019). We also note that Malvaez et al. (2015) studied the role of BLA glutamate signaling in an outcome-specific PIT procedure, employing two reinforcers and two instrumental response options. Here, we used single-outcome PIT, where a single instrumental response led to a single outcome. This is important, because a procedure using multiple reinforcers can stimulate different forms of learning (i.e., the formation of more detailed and sensory-specific neural representations of each reinforcer) compared to procedures using a single reinforcer (Corbit & Balleine, 2005; Blundell et al., 2001; Holland, 2004). Previous studies have also shown that different PIT paradigms can rely on different neural substrates. For example, the BLA can play different roles in outcome-specific versus general PIT (Corbit & Balleine, 2005). This being said, our results and those of Malvaez et al. (2015) combined suggest that glutamate signaling in the BLA is important across these PIT paradigms.

One consideration of the single-outcome PIT design used in our study is that both general and specific mechanisms might have contributed to the observed effects (Cartoni et al., 2016; Mahlberg et al., 2019). PIT designs that use a single outcome tend to be associated with general PIT, where the Pavlovian CS+ produces a general appetitive effect that can also potentiate instrumental responses for other outcomes (Cartoni et al., 2016). These single-outcome studies follow a general protocol almost identical to the present study, with the exception that here a second (inactive) lever was provided during instrumental training (Hall et al., 2001). Based on lesion studies by Blundell et al. (2001) and Corbit & Balleine (2005), the BLA is thought to mediate *specific transfer* rather than *general transfer*, so our findings may challenge this conclusion by demonstrating a role for BLA glutamate function in a PIT design biased towards general transfer. However, this is not conclusive as we do not have direct evidence that our design produces general transfer. A full transfer paradigm using two levers, each delivering a different outcome and three CSs (two paired with the outcomes delivered by the lever, and one paired with a third outcome) would be required to distinguish specific from general transfer effects (Corbit & Balleine, 2005; Cartoni et al., 2016). Nonetheless, our findings suggest that in animals with an intact BLA, glutamate-dependent processes within the BLA may indeed mediate general transfer, and this provides an avenue for further research.

### Limitations

Only male rats were studied in our experiments. Some studies suggest that female and male rats can differ in how they respond to reward-paired cues (Johnson et al., 2019; Nicolas et al., 2019; Zhou et al., 2014). Other studies suggest that female and male rats show similar cue-motivated behaviours (Collins et al., 2019). Nonetheless, there are limitations in generalizing findings obtained with males to females (Becker & Koob, 2016; Shansky, 2019; Shansky & Murphy, 2021), and so research should include animals of both sexes. In this regard, in ongoing work, we are examining potential similarities and differences between the sexes in PIT and in the effects of mGlu2/3 receptor activation.

### Conclusions

Activation of BLA mGlu_2/3_ receptors decreased the ability of a water-associated cue to goad both conditioned anticipation of water reward and water-seeking actions. We observed these effects under extinction conditions, in a pure conditioned incentive motivation paradigm that precluded the respective influences of primary or conditioned reinforcement on behaviour. The ability of increased BLA mGlu_2/3_ receptor activity to suppress the expression of conditioned incentive motivation did not involve general effects on motor behavior. We conclude that signaling via mGlu_2/3_ receptors within the BLA mediates the ability of reward cues to potentiate the incentive salience of their associated rewards.

## Acknowledgements

This work was supported by the National Science and Engineering Research Council of Canada (Grant #355923) and the Canada Foundation for Innovation to ANS (Grant #24326). ANS holds a salary award from the Fonds de la Recherche du Québec-Santé (Grant #28988). SYK was supported by a postdoctoral fellowship from the Fonds de la Recherche du Québec – Santé (Grant #270051). CG was supported by a Master’s scholarship (B1X) from the Fonds de recherche du Québec - Nature et technologies (Grant #275340). The authors gratefully acknowledge Rifka (Rebecca) C. Derman, Carrie R. Ferrario and Cameron Nobile for generously sharing equipment, computer programs and advice in setting up the behavioural tasks used in this work. The authors declare no conflicts of interest.

---

**Figure S.1.**
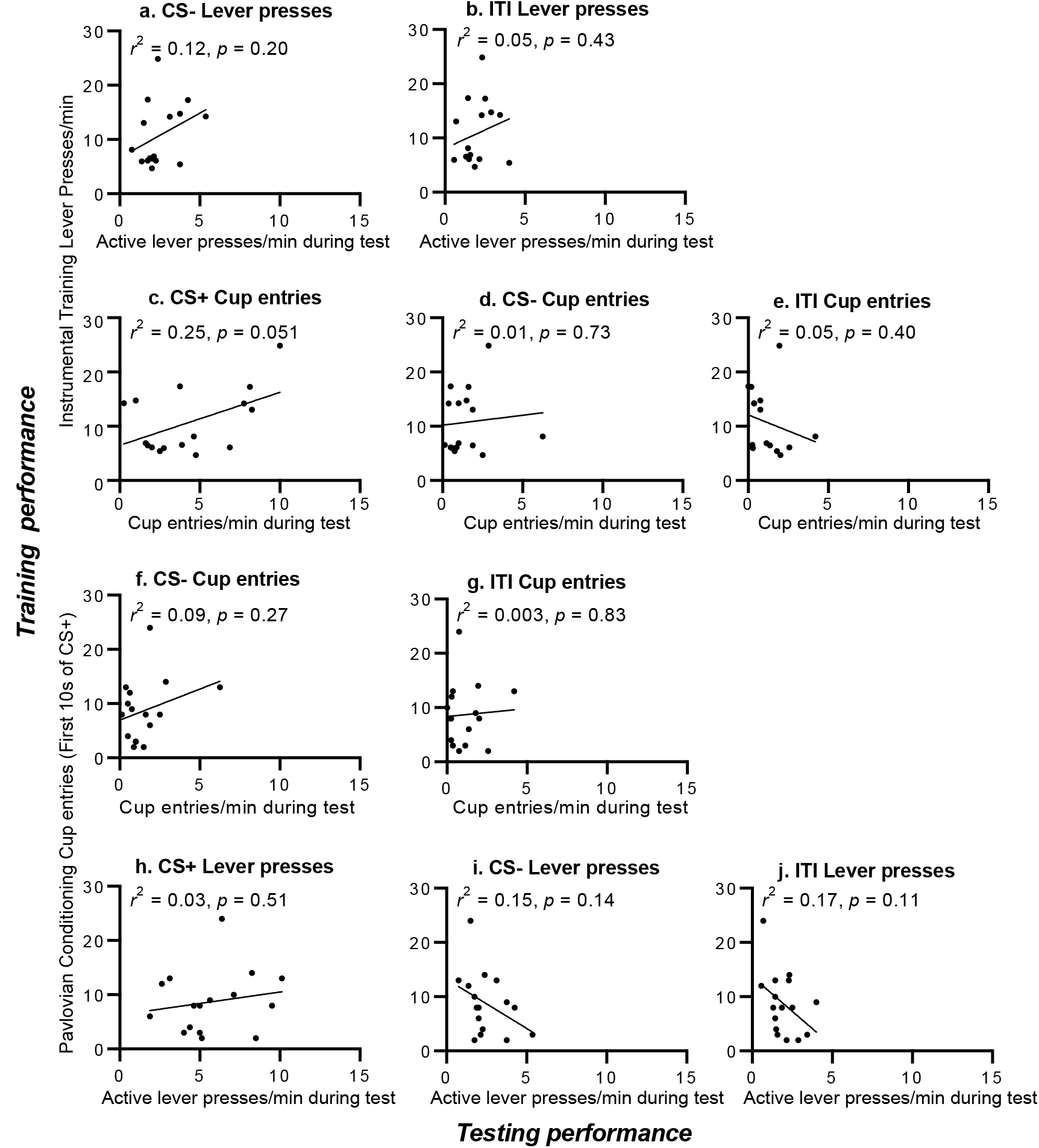
(a) More active lever pressing during the final instrumental training session did not significantly predict more active lever pressing at test during presentations of the CS-, (b) or the inter-trial interval. (c) Rates of active lever pressing during the final instrumental session also did not significantly predict water cup entries at test during the CS+, (d) during the CS- (e) or during the inter-trial interval (ITI). (f) Rates of water cup entries during the first 10 sec of CS+ presentation on the final Pavlovian session did not significantly predict water cup entries at test during the CS- (g) or inter-trial interval. (h) Water cup entries during the last Pavlovian conditioning session did not significantly predict active lever pressing at test during the CS+, (i) the CS- (j) or inter-trial interval. N = 16. Response rates during the CS+ and CS- are unadjusted for baseline (i.e., during the ITI). CS; conditioned stimulus.

